# Activation of group II metabotropic receptors attenuates cortical E-I imbalance in a 15q13.3 microdeletion mouse model

**DOI:** 10.1101/2020.09.17.301259

**Authors:** Marzieh Funk, Niklas Schuelert, Stefan Jaeger, Cornelia Dorner-Ciossek, Holger Rosenbrock, Volker Mack

## Abstract

Animal models reflecting human risk for schizophrenia are essential research tools for gaining further insight into the convergence of CNS pathology and clinical biomarkers. Amongst the variety of animal models that display schizophrenia-related neuronal network deficits, transgenic mice for rare and highly penetrant copy number variants (CNVs) provide a unique opportunity to study pathological correlates in models with strong construct validity. The Df(h15q13)/+ mouse model of the human 15q13.3 microdeletion CNV has been shown to mimic deficits in parvalbumin positive (PV+) interneuron and cortical network function. However, the corresponding changes in synapse density and activity within the medial prefrontal cortex (mPFC) have not been described. Using high-content immunofluorescence imaging, we have shown a reduced density of PV+ neurons and inhibitory synapses in the mPFC of Df(h15q13)/+ mice. We found that the reduced detection of PV+ synapses were accompanied by changes in spontaneous inhibitory and excitatory synaptic activity onto layer 2/3 pyramidal neurons. The aberrant cortical function was also evident in awake animals by a reduced high frequency auditory steady-state responses (ASSR), reliably monitored by EEG. Importantly, the imbalance of excitatory to inhibitory function could be attenuated on a cellular and cortical network level by activation of mGlu2/3 receptors, indicating the relevance of excessive excitatory transmission to the cortical network deficit in the Df(15q13)/+ mouse model. Our findings highlight the preclinical value of genetic risk and in particular CNV models such as the Df(15q13)/+ mice to investigate pathological network correlates of schizophrenia risk and to probe therapeutic opportunities based on clinically relevant biomarkers.

## Introduction

Schizophrenia is a complex neurodevelopmental disorder leading to a variety of devastating psychiatric symptoms^1^. The current treatment options for schizophrenia, which primarily target the dopaminergic system, are highly effective on positive symptoms but at best marginally improve negative symptoms or cognitive impairments^2, 3^. Therefore, identification of underlying mechanisms driving the symptoms in schizophrenia patients is required for the development of advanced and more effective treatments. Examination of postmortem prefrontal cortex (PFC) tissue from patients with schizophrenia has repeatedly demonstrated a significant reduction in the expression levels of the GABAergic interneuron marker parvalbumin (PV)^4, 5, 6^.

A major drawback in preclinical research has been the lack of informative animal models reflecting human risk and disease pathology^7^, for examining alterations in the activity of the complex interneuronal network in the PFC, which has been suggested to drive abnormalities in cortical network synchronization in patients with schizophrenia^8^. The identification of rare copy number variants (CNVs), which carry an increased risk for developing schizophrenia, provides a unique path towards translational studies in rodents^9–16^. The 15q13.3 microdeletion is a CNV resulting from a hemizygous deletion of approximately 1.5 Mb in the 15q13.3 locus, which significantly increases the risk of developing schizophrenia (odds ratio [OR] ~11)^9^ The same mutation has also been demonstrated to significantly increase the risk for developing autism, epilepsy and intellectual disability^17–20^. The generation of the respective mouse model Df(h15q13)/+ (15q13) by removing the syntenic genomic region encompassing 6 protein coding genes on chromosome 7 has enabled the exploration of pathological correlates and consequences. Two independently generated 15q13.3 microdeletion mouse models have been profiled and shown to demonstrate schizophrenia-related phenotypes such as cognitive impairments, social deficits and electrophysiological changes^21–23^.

Here we examined the connectivity of PV+ interneurons in the PFC of the 15q13 mouse model and identified changes in inhibitory synapse count and synaptic activity, correlating with an abnormal cortical E/I balance. Furthermore, we demonstrated that excessive excitatory transmission and deficits in fast cortical network processing were amenable to restoration using an mGluR2/3 agonist.

## Materials and Methods

### Animals

The Df(h15q13)/+ mouse was generated by Taconic Artemis (Köln, Germany) as previously described^21^. We used 10-11 week old 15q13 male mice for immunohistochemistry and in vivo EEG studies and 7-8 week old male mice for *in vitro* electrophysiology experiments (Taconic, USA). Animals were housed with a maximal of 4 per cage, with a minimum of 3 animals per cage. The animal facility was maintained under artificial lighting (12 h; light on at 6 am, light off at 6 pm), with both controlled ambient temperature and relative humidity. All experimental procedures were authorized by the Local Animal Care and Use Committee in accordance with local animal care guidelines, AAALAC regulations and the USDA Animal Welfare Act.

### Immunohistology

Immunohistological analysis was performed using slices from prefrontal cortex (Bregma 1.6 and 1.7) from 10 week old 15q13 mice and wild type littermates (N=6-8/group, 4 slices / animal). Mice were anesthetized with isoflurane and transcardially perfused with 0.1 M PBS (pH 7.4) followed by 4% paraformaldehyde. Brains were post fixed overnight in 4% paraformaldehyde and placed in 30% sucrose. Serial coronal sections (30μm thick) containing the prefrontal cortex were cut using a freezing stage sliding microtome (HM 450), collected in PBS (Invitrogen) and stored at 4°C. Immunofluorescent staining was performed on free-floating sections incubated with in 0.3% Triton in PBS for 20 minutes and then blocked with 10% normal goat serum in PBS for 45 min at room temperature. Sections were incubated with the following antibodies and lectin reagent: GAD67 (Mouse, Millipore, MAB5406,1:500); CamKII (rabbit, abcam, ab50202, 1:500); parvalbumin (guinea pig, Synaptic Systems, 195 004, 1:500); Gephyrin (Mouse, Synaptic Systems, 147 021, 1 in 500); biotin-conjugated Lectin Wisteria floribunda Agglutinin (WFA), (Sigma, L1516, 1:2000); NeuN (Mouse, Millipore, MAB377 1:500); Homer-1 (Rabbit, Synaptic Systems, 160002, 1:500); VGluT1 (Guinea pig, Synaptic Systems, 135304, 1:500). After a washing step with PBS for 45 minutes, the sections were incubated with the following secondary antibodies and streptavidin reagent for 2h at room temperature: goat anti-mouse IgG Alexa Fluor 488 (1:1000, ThermoFisher), goat anti-rabbit CF568 (1:1000, Sigma) and goat anti-guinea pig IgG Alexa Fluor 647 (1:1000, ThermoFisher) and Streptavidin Alexa Fluor 647 conjugate (ThermoFisher S32357). After another washing step with PBS/0.1% Triton for 1h, sections were mounted in 24-well glass bottom plates (Sensoplate, Greiner), air dried for 30 minutes and covered with aqueous mounting medium (Fluoroshield™ with DAPI, Sigma). Fluorescent images were taken and processed with an Opera Phenix (PerkinElmer) using the 63x objective in confocal mode.

### High content imaging analysis

For each brain section 330 visual fields were recorded, from which 24 fields were selected for analysis covering the pre-limbic and infralimbic cortex region according to the Allen Brain Atlas. Acapella Studio 4.1 software (PerkinElmer) was used to design a script for determining defined read-out parameters: Percent parvalbumin (PV), GAD67 and CamKII positive cells as well as PV bouton density (PV puncta per area). Invalid fields were excluded from the analysis due to insufficient nuclei count (below 20 nuclei per field /area205 μm x 205 μm). The DAPI staining was used to determine a nucleus mask. GAD67 positive cells were determined by calculating the GAD67 intensity within and a 6-7 μm ring region around the nucleus. For GAD67 positive cells the intensity ratio of nucleus/ring had to exceed a factor of 1.5. PV positive cells were determined analogously to GAD67 positive cells. PV positive puncta were determined by discriminating foreground objects (5-65 pixel = 0.2 μm2 to 2.6 μm2) with 1.3-fold higher intensities than the local background. The local background mask was determined as a ring region of 0.2 μm - 0.4 μm around individual PV puncta. The WFA intensity (perineuronal net abundance) was determined by creating a ring region around the PV cells (0-4 μm around PV soma) and calculating the mean WFA intensity within this ring region.

### Prefrontal cortex slice preparation

Mice were anesthetized with isoflurane and decapitated at the age of 7-8 weeks. After brain dissection, 250 μm thick coronal PFC slices were cut in an ice-cold oxygenated solution (87 mM NaCl, 2.5 mM KCl, 1.25 mM NaH2PO4, 7 mM MgCl2, 0.5 mM CaCl2, 25 mM NaHCO3, 25 mM glucose, 75 mM sucrose; 347 mOsmol at pH 7.4) using a vibratome (Leica VT1000S). Slices were stored in the same solution at 35 °C for 30 min and then transferred into recording ‘external’ solution at room temperature (124 mM NaCl, 3 mM KCl, 2.5 mM CaCl2, 1.3 mM MgSO4, 26 mM NaHCO3, 1.25 mM NaHPO4, 15 mM glucose; pH 7.4 when bubbled with 95% O2/5% CO2).

### Electrophysiology

Slices were placed in a submerged chamber on the stage of an upright microscope (Leica DM6 FS) and perfused at ~2 ml/min with recording solution (above). Layer 2/3 pyramidal neurons were visualized using epifluorescence illumination and infrared differential interference contrast illumination. Recording pipettes were pulled from thick-walled borosilicate glass tubing (1.5 mm outer diameter, 0.86 mm inner diameter, Harvard Apparatus). For whole-cell recording, pipettes were filled with internal solution (125mM CsMeSO3, 2mM CsCl, 10mM Na-HEPES, 5mM EGTA, 2mM MgCl2, and 4mM MgATP) and had a resistance of 5–10 MΩ. Currents were recorded at 22–26°C using an Axopatch-700B amplifier (Molecular Devices), filtered at 5 kHz (low-pass 8-pole Bessel filter) and sampled at 10 kHz.

Spontaneous inhibitory postsynaptic currents (sIPSCs) were recorded at 0 mV in the presence of 1 μM strychnine hydrochloride, 20 μM 6,7-dinitroquinoxaline-2,3-dione (DNQX) and 20 μM D-(-)-2-amino-5-phosphopentanoic acid (D-AP5) (Tocris Bioscience) to block glycine-, AMPA- and NMDA receptors, respectively. In all cases when tested sIPSCs were fully blocked by 20 μM SR-95531 (Tocris Bioscience). Spontaneous EPSCs (sEPSCs) were recorded at 70 mV after GABA_A_-, and glycine receptor were blocked by adding 20 μM SR-95531, and 1 μM strychnine (Ascent Scientific). To test the effect of mGluRII activation on spontaneous synaptic currents in pyramidal neurons of the PFC we measured EPSC and IPSC properties following 10 minute bath application of the mGluR2/3 agonist LY379268 (100nM)^24^.

### sIPSC and sEPSC analysis

Recordings were analysed using the NeuroMatic plugin (version 2.8, www.neuromatic.thinkrandom.com) in IgorPro 8 (Wavemetrics Inc., US) and were detected using a scaled template algorithm based on rising and decaying exponentials using a detection threshold of 5pA^25^.

### EEG electrode implantation

The technique used for the EEG recordings has been described in detail previously^26^. Briefly, for electrode implantation, animals were anaesthetized using isoflurane (2-3% in oxygen) and placed in a stereotaxic frame with a controlled heating blanket. Under sterile conditions, a longitudinal incision was made and the skin flaps were retracted with clamps. The surface of the scull was cleaned with hydrogen peroxide solution (5%) and dried off with an air puffer. Five small holes were drilled into the scull without damaging the dura mater. Coordinates for electrodes above the auditory cortex were AP −2.7 mm, and ML 4, and for the PFC were AP +1.5 mm, and ML 1 mm, relative to bregma. The hole for the reference electrode was positioned medially above the cerebellum at −5.7 mm AP. Then, 5 gold-plated steel screws each with a connector cable and a metal pin were implanted into the cranium superficially of the dura mater. A socket with the connector pins to attach the Neurologger (TSE systems) was implanted and fixed with dental cement. At the end of the surgery, animals received an injection of meloxicam, metacam 0.01 mg/kg i.p., and baitril s.c. over 5 days. After surgery, animals were maintained in individual cages and left to recover for 1 week.

### EEG recordings

The setup specifications, the different stimulation protocols and the recording procedures for the 40 Hz ASSR EEG experiments are described in detail in^26^. Briefly, auditory stimuli were generated with an audio generator and presented via a speaker system. An ASSR session consisted of a periodic train of single white noise clicks at a frequency of 40 Hz. Each train lasted 2 s with an interval of 10 s in between click trains. A total of 300 trains were presented. The intensity of the 40 Hz click train was adjusted to be 85.0 ± 1.0 dB. The mismatch negativity (MMN) protocol was described previously^26^.

Neurologger pre-amplifiers were attached to the implanted connector pins 30 minutes before starting the stimulation protocols. Subsequently, animals were individually placed in a novel open home cage in sound-attenuated recording boxes (MED Associates Inc.). The bottom of the cage was covered with a cotton tissue for rodent cages. To minimize the effects of stress, animals were given 30 min to acclimate to the apparatus prior to each recording session.

### EEG Data analysis

Data analysis was performed as described in^26^. The sampling rate of recording channels and trigger channels was set to 1000 Hz. The quality of EEG recordings was carefully checked and segments with artefacts were removed. Data were analysed using the Analyzer2 software package (BrainProducts GmbH, Munich, Germany). Data were segmented into 2.8 s epochs with 400 ms pre-trial period, 2 s stimulation period and 400 ms post-trial period. A Morlet wavelet analysis was performed for the 40 Hz frequency layer to measure mean power and inter-trial coherence (ITC) during the trial period. ASSR power was expressed as the accumulated power of the 40 Hz layer during the stimulation phase of 2 s. For the ITC, wavelet coefficients with complex numbers were computed and the phase-locking factor was determined by averaging the normalized phase synchronisation across trials for every time point and frequency.

### Compound administration

For the EEG recordings, the animals (11 weeks of age) were removed from their home cage and habituated for 15 minutes before recording the initial baseline. Following the initial baseline further recording started 15 min after compound administration. 15q13 mice and WT littermates received a single dose of 3 mg/kg LY379268 dissolved in saline injected s.c. at 10 weeks of age. Animals were randomly assigned to 3 groups containing equal numbers of 15q13 and WT mice that were recorded on 3 consecutive days.

### Data presentation and statistical analysis

Summary data are presented in the text as mean ± s.e.m. from N number of animals, as indicated. Comparisons involving two data sets were performed using a two-sided Welch two-sample t-test that does not assume equal variance. Comparison of EEG readouts between the WT and 15q13 cohort as well as differences between control-baseline and compound administration were analysed with a modification of the t-test of Satterthwaite for unequal variance of normally distributed sample groups. In the analysis, we do not assume a time effect.

## Results

### Diminished detection of PV^+^ cells and synapses in the PFC of 15q13 mice

We first examined whether we could detect alterations in the density and activity levels of PV^+^ in the 15q13 mice by immunolabeling sections of the mPFC with antibodies against markers for PV and Wisteria floribunda agglutinin (WFA), which is used for the detection of perineuronal proteins (PNNs) known for efficiently ensheathing PV cells^27^ (**Figure 1A-B**). Consistent with previous findings, we identified a reduction in the number of PV^+^ interneurons in the prelimbic (PL) and infralimbic region (IL) of the mPFC of 15q13 mice (t=_5.51_ P= 1.31^-06^ and t=_3.49_ P= 0.0010 respectively), along with a significant reduction in the somatic PV intensity in the IL (t=_2.22_ P=0.032) (**Figure 1C**). Additionally, we detected a reduction in the intensity of WFA-labelling in the PL and IL of 15q13 mice (t=_4.812_, P=8.58^E-06^ PL and t=_3.45_ P=0.001 IL) without significant alteration in the fraction of PV+ cells surrounded by PNNs (t=_0.771_ P=0.440 PL and t=_1.12_ P=0.269 IL) (**Figure 1D**). To determine whether the reduced density of PV^+^ cells in the PFC of the 15q13 mice was the result of general neuronal loss, we assessed the number of NeuN-positive neurons. We detected no significant change in the number of labelled neurons in the PFC of 15q13 mice (t=_1.360_ P= 0.180 and t=_1.425_ and P=0.161 respectively), suggesting that reduced PV protein expression may cause the lower density of detected PV+ cells (**Supplementary Figure 1 A-B**).

**Figure. 1.**
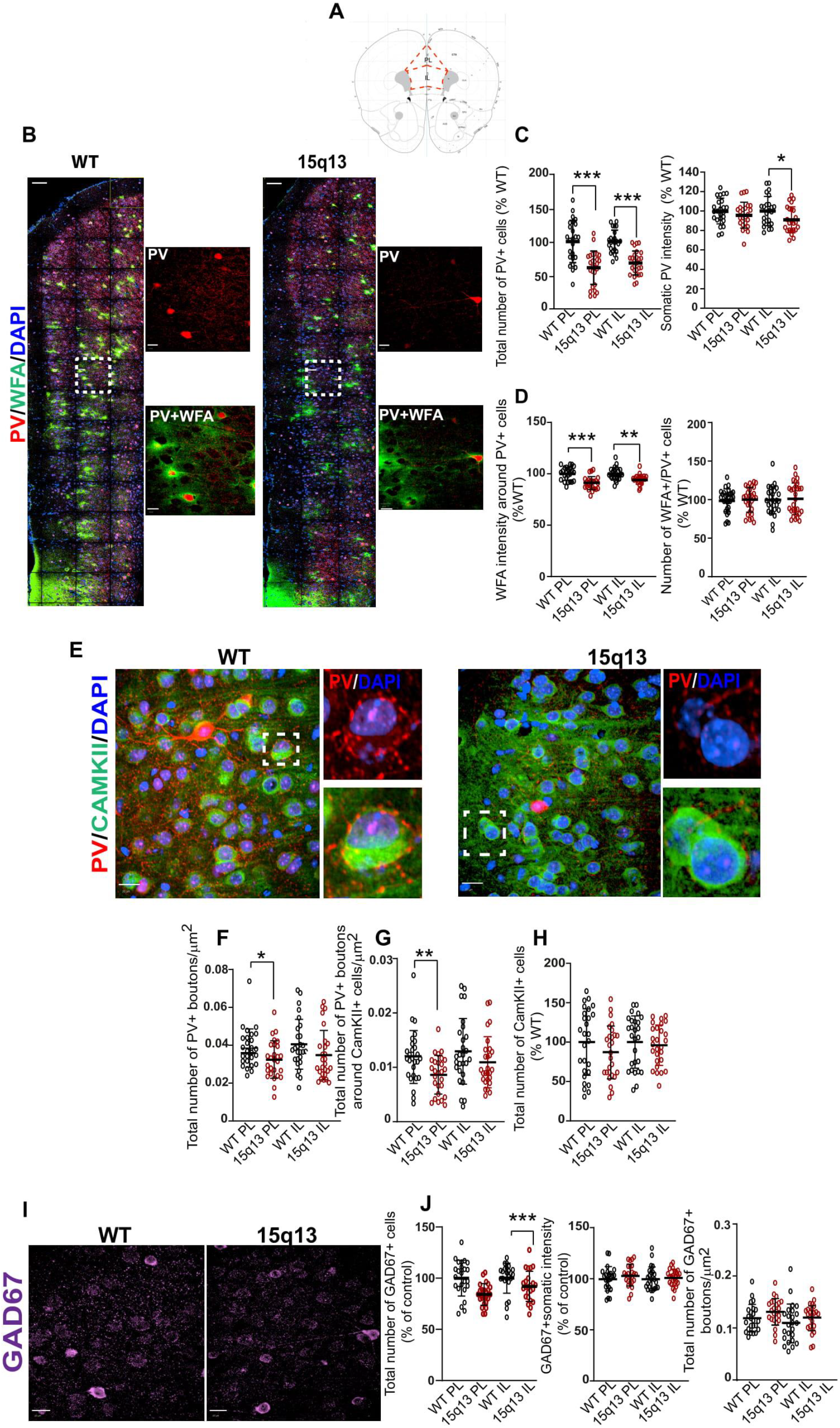
Reduced number of inhibitory cells and synapses in the prefrontal cortex of the 15q13 mouse model. **A** Schematic diagram of the location of the prelimbic (PL) and infralimbic region (IL) of the mouse prefrontal cortex. **B** Representative confocal images from wild-type (WT) and 15q13 mice (15q13) prefrontal cortex labelled with WFA and PV antibodies. Scale bar 100μm. **C-D** Summary graphs showing the effect of 15q13.3 microdeletion in PL and IL on the number of PV+ cells, intensity of the somatic PV signal together with the intensity of PNNs ensheathing PV+ cells (n = 24 sections from 6 WT and 6 15q13 mice). Graphs show mean ± s.e.m. * P < 0.05, ** P < 0.01, *** P < 0.001 (unpaired Welch two-sample t test). **E** Representative confocal images from wild-type (WT) and 15q13 mice (15q13) prefrontal cortex labelled for PV (red), CamKII (green), and DAPI (blue). Scale bar 20μm. Selected regions of the PL (white box) are enlarged on the right. Scale bar 20μm. **F-H** and the number of PV+ boutons in total and surrounding CamKII+ neurons, as well as the number of CamKII+ cells (n = 32 sections from 8 WT and 8 15q13 mice). **I** Representative confocal images of WT and 15q13 PL stained for GAD67 (magenta). Scale bar 20μm **J** Pooled data showing the reduction in the number of GAD67+ cells in the IL of the 15q13 mice, without significant alteration in somatic GAD67 expression and number of boutons in the PFC. (n = 32 sections from 8 WT and 8 15q13 mice). Graphs show the mean ± s.e.m. * P < 0.05, ** P < 0.01, *** P < 0.001 (unpaired Welch two-sample t test).

To study whether the reduced number of PV+ cells was accompanied by diminished number of respective synapses and inhibitory drive onto excitatory neurons we quantified the number of PV+ boutons and their density on the soma of CamKII+ excitatory neurons in the mPFC (**Figure 1E**). We identified a significant decrease in the total number of PV+ boutons (t=_2.21_ P= 0.0316) and number of PV+ boutons contacting CamKII+ neurons in the PL (t=_2.830_ P= 0.007) without significant decrease in the IL of 15q13 mice (t=_1.610_ P=0.113, t=_1.372_ P= 0.176 respectively) (**Figure 1F-G**). In contrast to the reduced detection of PV+ cells in the mPFC, we did not detected a significant alteration in the number of CamKII+ excitatory neurons in the PFC of the 15q13 mouse model was not changed (t=_1.46_ P=0.15) (**Figure 1H**). Together, these results indicate lower activity of PV^+^ interneurons and reduced inhibitory input onto excitatory neurons in the mPFC of 15q13 mice.

Previous studies have demonstrated a reduced expression of the 67-kDa isoform of glutamic acid decarboxylase (GAD67), an enzyme for γ-aminobutyric acid (GABA) synthesis in the mPFC of schizophrenia patients^28^. Using high content imaging, we explored whether the expression of GAD67 were affected in the mPFC of the 15q13 mouse model (**Figure 1I**). We detected a lower number of GAD67+ cells in the IL (t=_1.904_ P= 0.063 PL and t=_3.81_ P=0.0004 IL), without significant changes in intensity (t=_1.095_ P= 0.279 PL and t=_0.415_ P= 0.681 IL) or number of GAD67+ boutons in the mPFC of 15q13 mice (t=_1.194_ P= 0.238 PL and t=_1.646_ P= 0.106 IL) (**Figure 1J**). Together, our results revealed further convergence on markers of abnormal interneuron function in the mPFC in the 15q13 mouse model and schizophrenia patients.

In addition, we examined whether the changes in PV+ immune detection in the PFC were accompanied by alterations in markers for excitatory synapses. We labelled PFC sections for Homer1, VGluT1 and CamKII (**Supplementary Figure 2**) but did not detect significant changes in the number of excitatory synapses in the PFC of 15q13 mice (t=_1.628_ P= 0.110 PL, t=0.5057 P= 0.616 IL) (**Supplementary Figure 2B**). This provides additional support for the crucial impact of interneuron activity as potential driver for aberrant cortical network function in the 15q13 mouse model^29^.

### Reduced inhibitory and increased excitatory synaptic currents in layer 2/3 pyramidal neurons of the 15q13 mice

We explored whether the reduction in PV+ expression correlated with functional deficits in the PFC of 15q13 mice by measuring the spontaneous inhibitory currents (sIPSCs) in layer 2/3 pyramidal cells using whole-cell electrophysiological recordings (**Figure 2A-C**). We identified a significant reduction in sIPSC frequency (WT 3.66 ± 0.92 Hz to 1.38 ± 0.36Hz 15q13 t=_2.42_ P=0.0268; **Figure 2D**) without alteration in average sIPSC amplitude (WT 9.82 ± 3.30pA to 8.78 ± 0.903pA 15q13 t=_0.32_ P= 0.75; **Figure 2E**), consistent with a reduced expression of PV in the PFC of the 15q13 mouse model. The reduction in the inhibitory drive onto cortical pyramidal neurons has been linked to a hyperglutamatergic state in the PFC of patients with schizophrenia^30^. We therefore recorded spontaneous excitatory currents (sEPSCs) from layer 2/3 pyramidal neurons (**Figure 2F-G**), and identified a significant increase in sEPSC frequency (WT 7.75 ± 1.46Hz to 21.79 ± 4.40Hz 15q13 t=_2.38_ P=0.0312; **Figure 2H**), without a change in amplitude (WT 8.68 ± 1.831pA to 10.90 ± 1.21pA 15q13 t=1.05 P=0.3098; **Figure 2I**). Our results provide evidence for an impaired inhibitory control of cortical pyramidal cells and elevated excitatory activity in the PFC of the 15q13 mouse model.

**Figure. 2.**
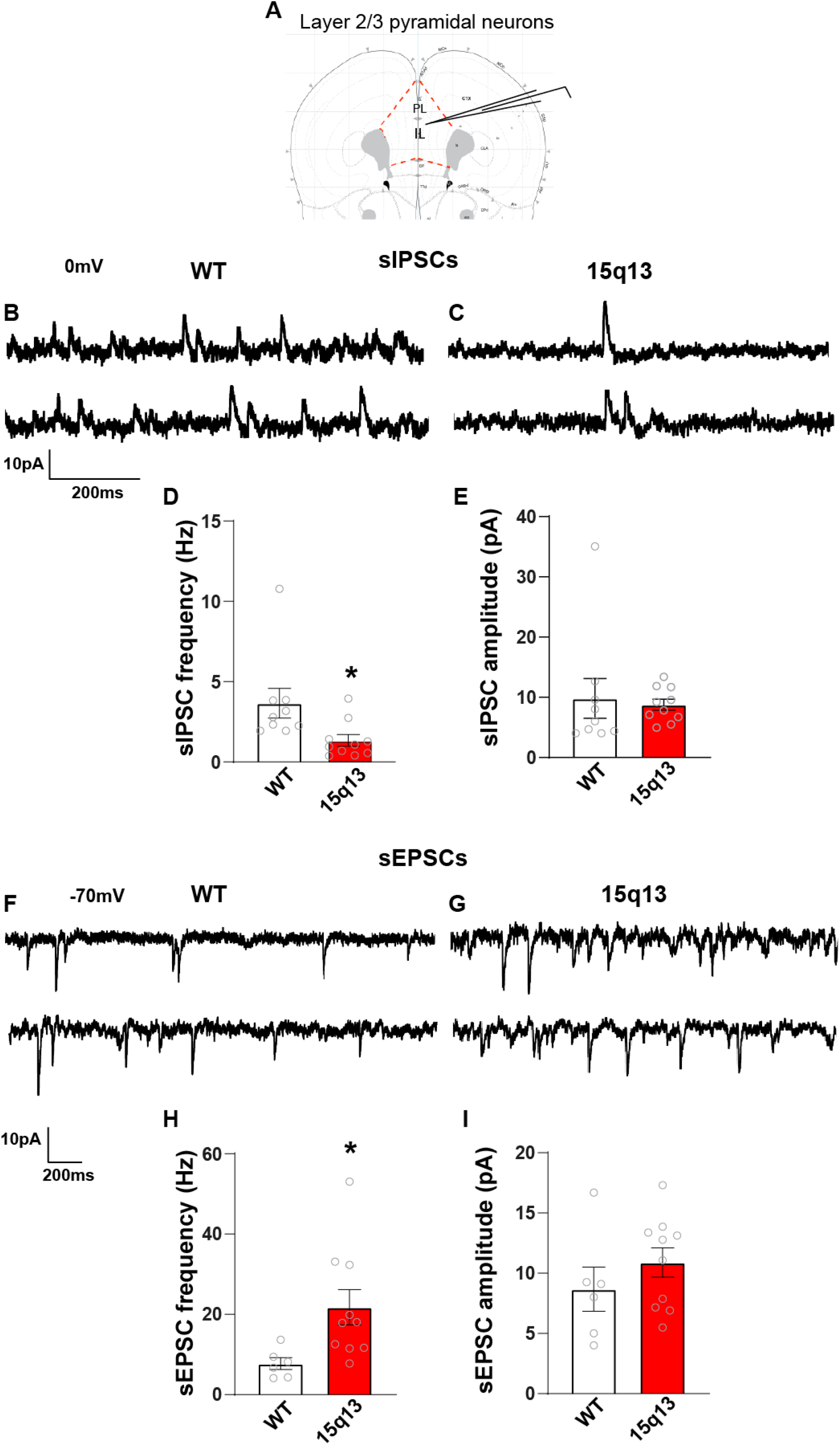
Imbalance in excitatory and inhibitory basal transmission in layer 2/3 pyramidal neurons in the mPFC of the 15q13 mouse model. **A** Schematic diagram of the location for the electrophysiological recordings from layer 2/3 mPFC. **B-C** Representative traces of sIPSCs recorded at 0mV from layer 2/3 pyramidal neurons from WT and 15q13 mice. **D-E** Pooled data showing reduced sIPSC frequency with no change in the amplitude in the 15q13 mouse model (n = 9 cells from n = 8 WT mice and 10 cells from 9 15q13 mice). **F-G** Representative traces of sEPSCs from WT and 15q13 layer 2/3 pyramidal neurons (−70mV). **H-I** Pooled data showing increased sEPSC frequency in layer 2/3 pyramidal neurons from 15q13 mice with no change in amplitude. (n = 6 cells from 6 WT mice and n = 10 cells from 8 15q13 mice). All graphs are presented as mean ± s.e.m. * P < 0.05, ** P < 0.01, *** P < 0.001 (unpaired Welch two-sample t test).

### Attenuating presynaptic glutamate release normalised synaptic hyperexcitation in layer 2/3 pyramidal neurons of the 15q13 mice

The increase in sEPSC frequency indicated excessive glutamate transmission as an integral part of the cortical pathology in 15q13 mice, which may affect E/I balance and impair neuronal network function. We probed this hypothesis in acute brain slices by pharmacological modulation of excitatory transmission. The activation of mGlu2/3 receptors have previously been shown to attenuate synaptic release of glutamate, thereby downregulating network excitability^24, 31, 32^. Following bath application of the mGluR2/3 agonist LY379268 (100 nM for 10 min) (**Figure 3A**), sEPSCs frequency in pyramidal neurons from layer 2/3 pyramidal neurons were significantly reduced (baseline 17.58Hz ± 2.14 to 4.80Hz ± 0.86 LY379268 t=_5.53_ P=0.0006; **Figure 3B**) with no alteration in amplitude (baseline 7.78pA ± 1.84 to 7.17pA ± 1.26 LY379268 t=_0.27_ P=0.790; **Figure 3C**). Previous studies have demonstrated that the activation of mGlu2/3 receptors can also modulate the release of GABA and thereby inhibitory transmission^33^. We recorded sIPSCs following bath application of LY379268 (**Figure 3D**), and in contrast to the normalisation of the sEPSC frequency, LY379268 did not significantly restore inhibitory synaptic transmission in the 15q13 mouse model (baseline 1.16Hz ± 0.380 to 2.05Hz ± 0.721 t=_1.659_ P=0.135; **Figure 3E-F**). Taken together, these results confirm that attenuation of synaptic glutamate release by the mGluR2/3 agonist LY379268 can revert the hyperexcitation of pyramidal neurons in the PFC of 15q13 mice.

**Figure. 3.**
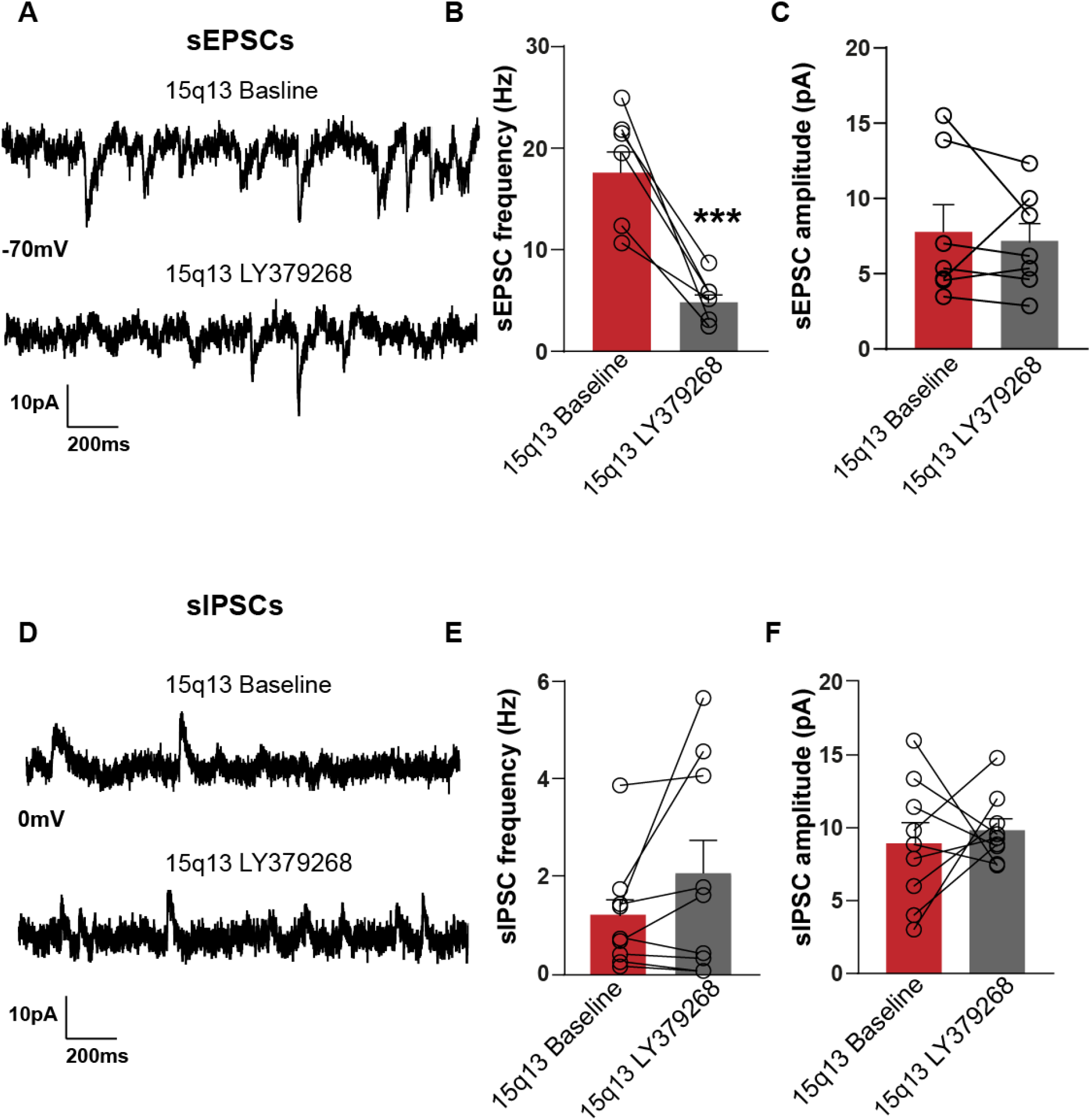
Activation of mGluR2/3 attenuates hyperexcitation in layer 2/3 pyramidal neurons of the 15q13 mouse model. **A** Representative sEPSC traces in voltage clamp mode (−70mV) during baseline and following bath application of the mGluR2/3 agonist LY379268 (100 nM) recorded from layer 2/3 pyramidal neurons in the 15q13 mouse model. **B** Summary plot showing the attenuation of sEPSC frequency of layer 2/3 pyramidal neurons from 15q13 mice in response to LY379268 (100nM) (n = 7 cells from 7 15q13 mice). **C** Pooled data showing the lack of effect of LY379268 on the amplitude of sEPSCs in layer 2/3 pyramidal neurons. **D** Representative sIPSC traces (0mV) from layer 2/3 pyramidal neurons from baseline and following 100nM bath application of LY379268 (n = 9 cells from 8 15q13 mice). **E-F** Pooled data showing no significant effect of LY379268 on the sIPSC frequency and amplitude in layer 2/3 pyramidal neurons. All graphs are presented as mean ± s.e.m. * P < 0.05, ** P < 0.01, *** P < 0.001 (unpaired Welch two-sample t tests).

### Reversion of cortical processing deficits in 15q13 mice by an mGluR2/3 agonist

As a next step we aimed to evaluate changes in cortical sensory processing in 15q13 mice, using multiple auditory stimulation protocols for recording evoked EEG patterns in vivo (**see Methods**). Recording electrodes were fixed to measure epidural EEG signals from auditory and prefrontal cortex regions (**Figure 4A-B**). As reported by others, we determined changes in cortical high-frequency processing as deficits in the 40 Hz ASSR power (**Figure 4C, D and Table 1**)^21, 34^ and inter-trial phase coherence (ITC) in 15q13 mice (**Figure 4F, G and Table 1**). Other schizophrenia-relevant readouts, such as mismatch negativity (MMN) or basal cortical oscillations in the gamma frequency range, were not significantly altered (**Table 1**). Since the 40Hz cortical ASSR deficit represents a readout for impaired PV+ interneuron function and resulting E-I imbalance we wanted to assess how suppression of glutamatergic transmission may affect the marker of fast cortical processing in 15q13 mice. Importantly, administered of the mGluR2/3 agonist LY379268 at 3 mg/kg, previously shown to reduced glutamate release *in-vivo*^35, 36^, led to an acute normalization of ASSR power in the PFC and auditory cortex (AC), as well as ITC in the PFC in 15q13 mice (**Figure 4E, H and Table 1**). Not surprisingly, the compound did also augment parameters of cortical network processing in WT mice (**Table 1**). Vehicle administration did not significantly affect respective EEG pattern in either group (**Supplementary Figure 3**). Even though the compound has shown activity on various EEG readouts regardless of the genotype, our data support a beneficial effect of mGluR2/3 activation on cortical network function based to an emerging EEG biomarker for schizophrenia.

**Figure. 4.**
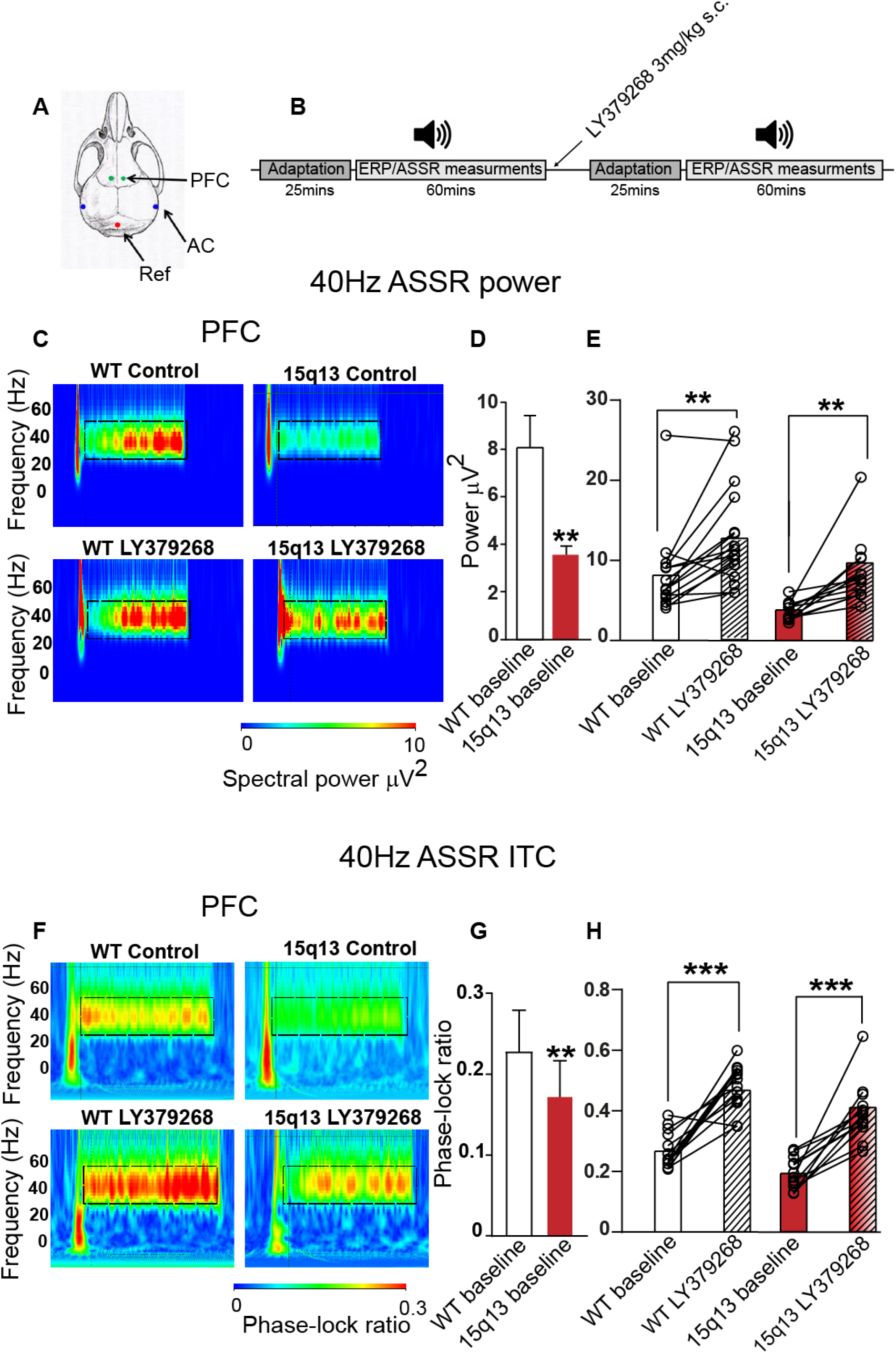
The mGluR2/3 agonist LY379268 normalises deficits in auditory state response (ASSR) in the 15q13 mouse model. **A** Image representing location of EEG electrodes above the prefrontal cortex (PFC) and auditory cortex (AC). **B** Timeline showing the experimental design. **C** Representative time frequency plots from an individual WT (left images) and 15q13 (right images) mice showing 40 Hz ASSR power in response to a given 40-Hz click trains (85 dB intensity, 2 sec duration) at baseline and following a single s.c injection of 3mg/kg of LY379268. **D** Summary graph showing the significant reduction in the average ASSR power in the mPFC from WT and 15q13 mice (n = 16 WT and 14 15q13 mice). **E** Pooled ASSR data from WT and 15q13 mice showing the significant increase in ASSR power following s.c. administration of 3mg/kg of LY379268 (n = 15 WT and 12 15q13 mice). **F** Representative examples of spectrograms from the PFC of WT and 15q13 mice demonstrating the intra trial coherence (ITC) in response to a 40Hz steady state tone. **G** Summary graph of the ITC in the PFC of the 15q13 mice (n = 14 WT and 14 15q13 mice). **H** Pooled data showing the significant increase in ITC in the PFC of 15q13 mice following administration of LY379268 (n = 15 WT and 12 15q13 mice). Asterisks denote the significant difference between baseline power and ITC from WT and 15q13 mice and the difference between baseline and following administration of LY379268. All graphs are presented as mean ± s.e.m * P < 0.05, ** P < 0.01, *** P < 0.001 (Based on one-sample t-test for paired observations).

**Table 1.**
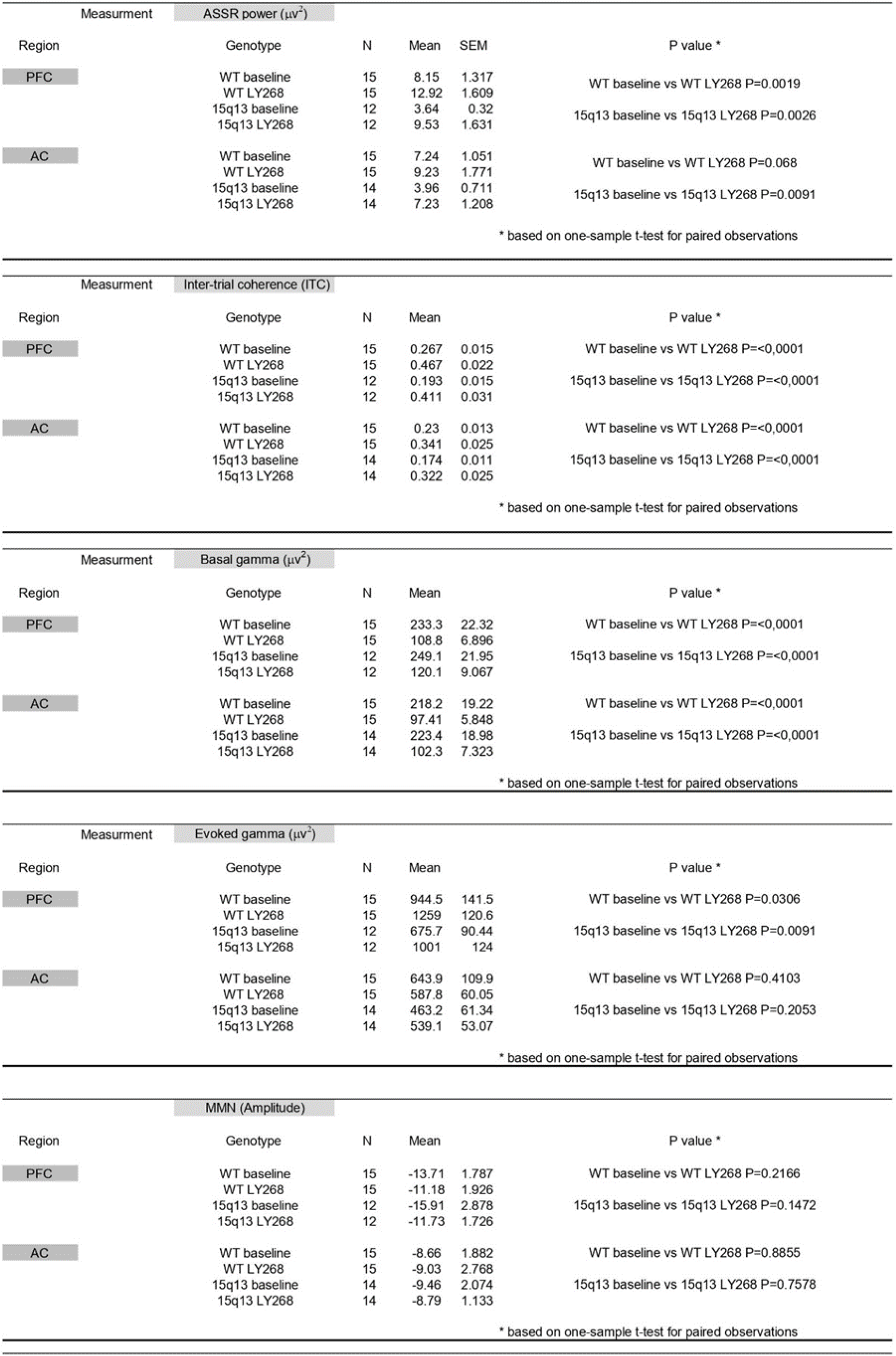
LY379268 significantly alters the 40Hz auditory state response and Inter-trial coherence (ITC) in the 15q13 mouse model. N indicates the number of WT and 15q13 animals used for the EEG measurements. Systemic administration of LY379268 significantly increased the basal and evoked gamma in the 15q13 and wild type control mice.

## Discussion

The reduced activity of fast-spiking PV+ interneurons in the cortex of 15q13 mice has been suggested to mediate deficits in cortical network synchronization^29^. However, it remained unclear how changes in the activity of interneurons may contribute to the impairment in cortical E/I balance. In this study, we revealed a decrease in PV+ synapses in the 15q13 model in conjunction with increased excitation of PFC pyramidal neurons. We demonstrated that pharmacological attenuation of glutamate transmission normalized cortical hyper-excitation and augmented cortical processing in 15q13 mice leading to a reversion of apparent gamma-band ASSR deficits. Our data emphasize the value of the 15q13 mouse model for the exploration of pathological and therapeutic mechanisms based on functional biomarkers associated with schizophrenia.^37^.

The activation of fast-spiking PV+ basket cells is critical for the regulation of cortical excitation, which is achieved via the dense innervation of the soma, dendrites, and axon initial segment of pyramidal neurons^38–40^. The reduction in PV+ cell-mediated inhibition in the PFC has been proposed to be a major cause for altered cortical network activity in patients with schizophrenia^41, 42^. Similar to pathological findings in schizophrenia^43^, we also showed a robust decrease in the number of PV+ cells and synapses together with a significant decline in inhibitory transmission onto pyramidal cells in the mPFC of the 15q13 mouse model.

Our observation of increased cortical excitatory synaptic activity in 15q13 mice has also been described for other models of schizophrenia risk, including the dysbindin-1 mutant mice (Dys-MT)^44^, Poly (I:C) mice^45^ and a Disc1 mouse model^46^. Importantly, elevated glutamate levels have been detected in the PFC of schizophrenia patients and linked to positive as well as negative symptoms^47, 48^. Additional support for the role of altered cortical glutamate drive in psychosis and schizophrenia comes from studies with systemic administration of the N-methyl-d-aspartate (NMDA) receptor antagonist ketamine, which results in increased glutamate efflux in the PFC together with perceptual abnormalities^49, 50^ as a potential consequence of disinhibition of PV+ interneurons^51^.

Various studies have demonstrated that specific moderation of excessive cortical glutamate transmission can be achieved by activation of mGlu2/3 receptors^52–54^. In our study, acute administration of the mGluR2/3 agonist LY379268 efficiently normalized glutamatergic transmission without a significant effect on inhibitory currents in PFC pyramidal neurons from 15q13 mice. Interestingly, we did not observe depression of inhibitory synaptic currents as detected previously^33^, which could be attributed to the differences in network connectivity and receptor distribution between the cochlear nucleus and the PFC.

In line with a reduction of excitatory transmission, we demonstrated a robust effect of LY379268 on cortical EEG responses, including the reversion of cortical gamma oscillation deficits in 15q13 mice. Cortical gamma oscillations have been shown to be dependent on synaptic connections between PV+ interneurons and pyramidal neurons^55, 56^, which are linked to a wide range of perceptual and cognitive processes believed to be crucial to functional disruptions in schizophrenia.

The 40-Hz auditory steady-state responses (ASSRs) represents a promising biomarker for monitoring high frequency network synchrony in schizophrenia patients^57–59^ and in subjects at ultra-high risk for developing schizophrenia^60^. Importantly, ASSR has also been correlated with a range of cognitive functions such as working memory, attention, reasoning and problem solving^60–65^ and may reflect neuronal network mechanisms of cognitive processing. Additionally, ASSR can be monitored in humans and rodents; therefore, it can be utilized as a translational cross-species biomarker for studying risk, pathological mechanisms and novel therapeutic strategies. Furthermore, changes in 40-Hz ASSRs have also been detected in other psychiatric disorders and relatives of schizophrenia patients, suggesting that the marker may indicate a converging trajectory of neuropsychiatric risk and provide insight into disease pathology^58^.

The 15q13 mice displayed particular deficits in cortical ASSR, which were recovered upon acute administration of the mGluR2/3 agonist LY379268. Activation of the mGlu2/3 receptors augmented the impaired 40Hz ASSR power and ITC and thereby emphasized the impact of an excessive glutamatergic activity on network dysfunction in the 15q13 mouse model. It should be noted that the activation of mGluR2/3 by LY379268 had a broad effect on multiple EEG readouts monitoring basal or evoked oscillations in wild type and 15q13.3 microdeletion mice. The effects on basal oscillations across multiple power bands and auditory evoked potentials in rats have previously been reported^66, 67^ and could be attributed to the general effect of mGluR2/3 activation on glutamate transmission.

As mentioned above, studies in humans have indicated a beneficial effect of mGluR2/3 agonists on ketamine-induced cognitive deficits^68^ and recently confirmed the normalization of cortical hyper-activity by magnetic resonance imaging^69^.

In conclusion, we provided further insights into cortical network changes contributing to the impaired E/I balance in a 15q13.3 microdeletion mouse model reflecting genetic risk for schizophrenia. We demonstrated that cortical hyper-excitation, possibly because of insufficient inhibitory control by PV+ interneurons, contributes to the impairment of auditory evoked gamma oscillations. The restoration of the ASSR deficits by activating mGlu2/3 receptors underscores the value of the 15q13 mouse model to monitor therapeutic mechanisms based on a translational biomarker of schizophrenia and once more indicates the potential clinical benefit of lowering excessive glutamatergic transmission.

## Funding and Disclosure

This work was supported by Boehringer Ingelheim. The authors have nothing to disclose.

## Acknowledgments

We thank Klaus Bornemann, Matthias Heil, Bernd Igl, Monika Bruening, Scott Hobson and Dennis Kaetzel for their help and useful discussions.

## Author contributions

MZ, NS, HR, CDC and VM designed the experiments. MZ performed the in-vitro electrophysiology and analysis. NS performed the in-vivo EEG and analysis. MZ carried out the immunohistochemistry and SJ analysed the data. MZ, CDC and VM wrote the manuscript.

---

**Supplementary Figure 1.**
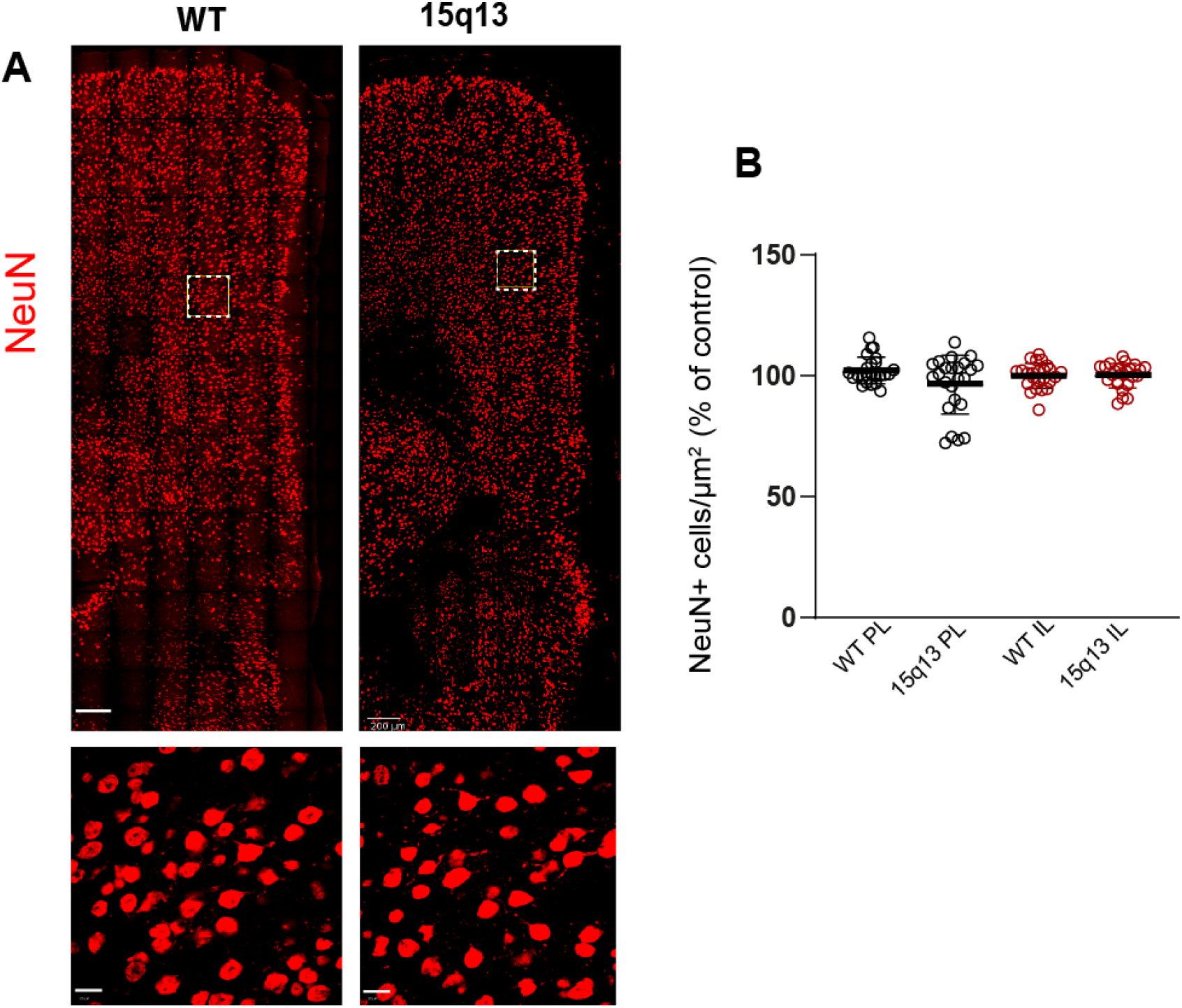
No alteration in the number of NeuN+ cells in the PFC of the 15q13 mouse model. **A** Representative confocal images from wild-type (WT) and 15q13 mice (15q13) prefrontal cortex labelled for NeuN. Scale bar 200μm. Selected regions of the PL (white box) are enlarged below. Scale bar 20 μm. **B** Pooled data showing the total number of NeuN+ cells in the PL and IL in the PFC of the 15q13 mouse model. (n = 24 sections from 6 WT and 6 15q13 mice). Graphs show mean ± s.e.m.

**Supplementary Figure 2.**
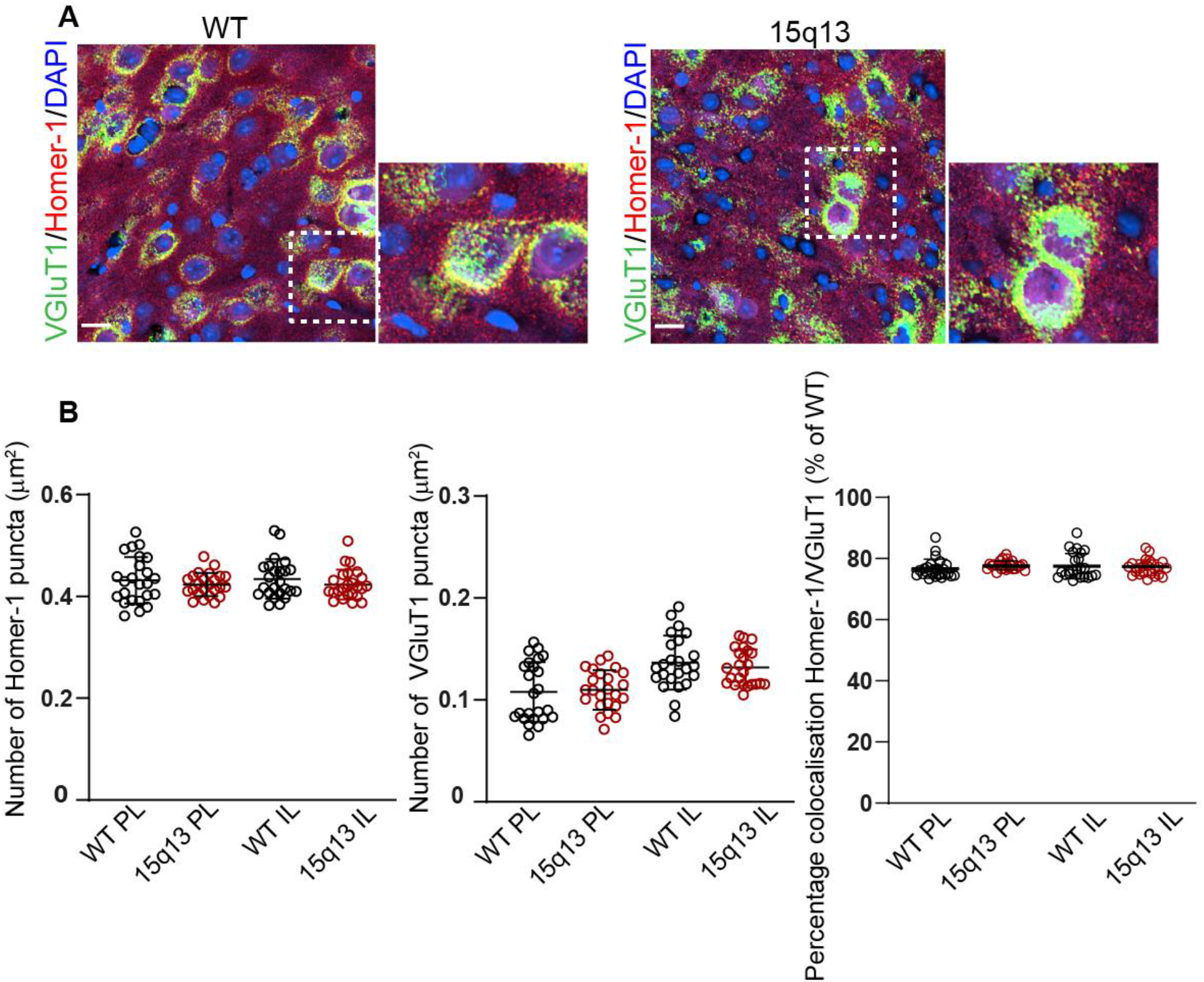
No significant alteration in the number of excitatory synapses in the PFC of the 15q13 mouse model. **A**. Representative images from wild-type (WT) and 15q13 mice (15q13) labelled with the excitatory synaptic markers: VGluT1 (green) Homer1 (red) and DAPI (blue). **B** Pooled data showing no significant reduction in the number of Homer-1 and VGluT1 in the PL and IL in addition to no alteration in the percentage of colocalisation of Homer1 and VGluT1 in the PFC of the 15q13 mouse model. (n = 24 sections from 6 WT and 6 15q13 mice). Graphs show mean ± s.e.m. * P < 0.05, ** P < 0.01, *** P < 0.001 (unpaired Welch two-sample t test).

**Supplementary Figure 3.**
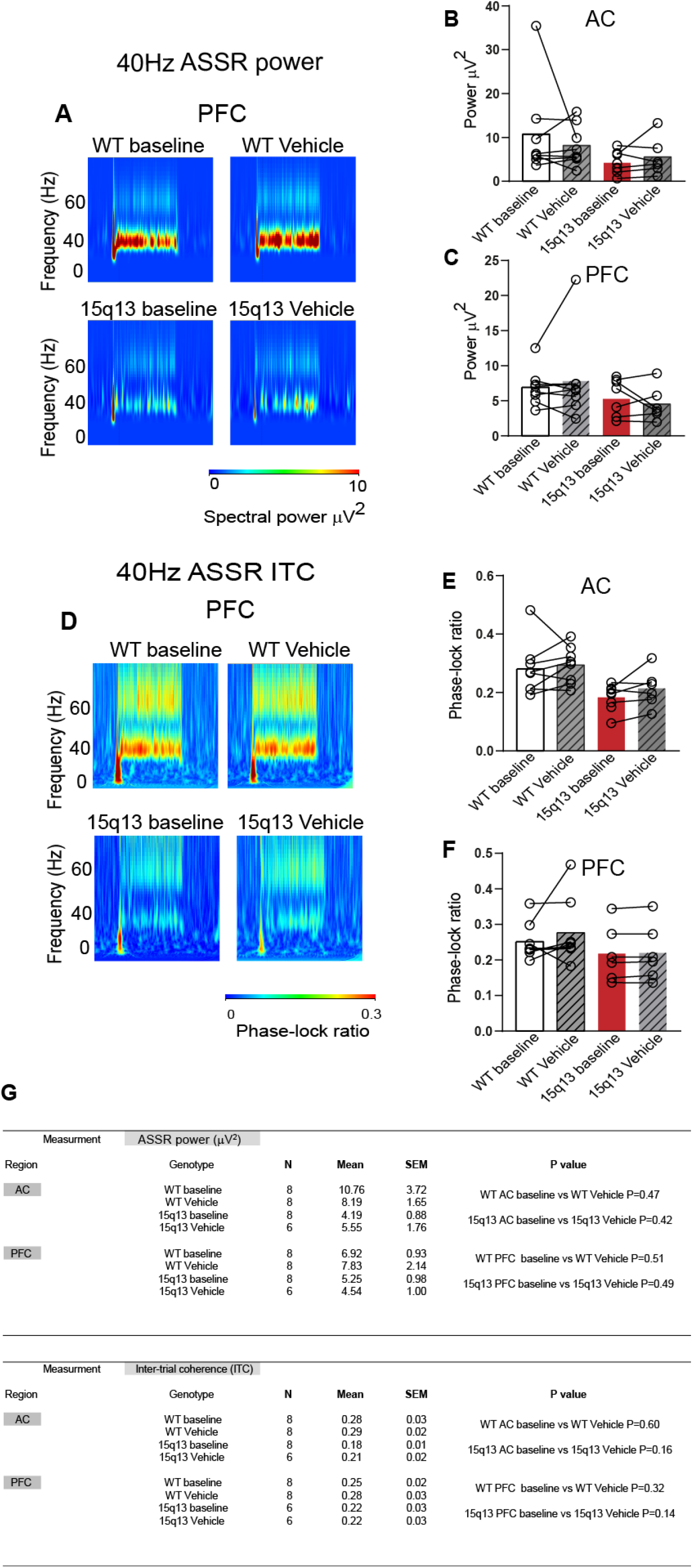
Vehicle administration of saline does not significantly alter the auditory state response (ASSR) in the 15q13 mouse model. **A**. Representative time frequency plots from an individual WT (left images) and 15q13 (right images) mice showing 40 Hz ASSR power in response in the auditory (AC) and the prefrontal cortex (PFC) to a given 40-Hz click trains (85 dB intensity, 2 sec duration) at baseline and following a saline injection (s.c). **B-C** Pooled ASSR power data from WT and 15q13 mice showing no significant change in the 40Hz ASSR power following s.c. administration of saline (n = 8 WT and 8 15q13 mice for AC and n = 8 WT and 6 15q13 mice for PFC). **D** Representative time frequency plots from an individual WT (left images) and 15q13 (right images) mice showing 40 Hz ASSR ITC in response in the auditory (AC) and the prefrontal cortex (PFC) to a given 40-Hz click train (85 dB intensity, 2 sec duration) at baseline and following a saline injection (s.c). **E-F** Pooled ASSR ITC data from WT and 15q13 mice showing no significant change in the 40Hz ASSR ITC following s.c. administration of saline (n = 8 WT and 8 15q13 mice for AC and n = 8 WT and 6 15q13 mice for PFC). **G** Statistical analysis table examining the effect of administration of saline (s.c) on the 40Hz auditory state response and Inter-trial coherence (ITC) in the 15q13 mouse model. N indicates the number of WT and 15q13 animals used for the EEG measurements. (Based on one-sample t-test for paired observations).

## Notes

### Competing Interest Statement

The authors have declared no competing interest.

